# Automatic Traits Extraction and Fitting for Field High-throughput Phenotyping Systems

**DOI:** 10.1101/2020.09.09.289769

**Authors:** Xingche Guo, Yumou Qiu, Dan Nettleton, Cheng-Ting Yeh, Zihao Zheng, Stefan Hey, Patrick S. Schnable

## Abstract

High-throughput phenotyping is a modern technology to measure plant traits efficiently and in large scale by imaging systems over the whole growth season. Those images provide rich data for statistical analysis of plant phenotypes. We propose a pipeline to extract and analyze the plant traits for field phenotyping systems. The proposed pipeline include the following main steps: plant segmentation from field images, automatic calculation of plant traits from the segmented images, and functional curve fitting for the extracted traits. To deal with the challenging problem of plant segmentation for field images, we propose a novel approach on image pixel classification by transform domain neural network models, which utilizes plant pixels from greenhouse images to train a segmentation model for field images. Our results show the proposed procedure is able to accurately extract plant heights and is more stable than results from Amazon Turks, who manually measure plant heights from original images.

## Introduction

High-throughput phenotyping is a new technology that takes images for hundred and thousands of plants simultaneous and continuously during their whole growth stages. It is constructed to improve the classical labor-intensive and inefficient hand-measured approach for collecting plant traits. Substantial advancements have been made by engineers to enable the large-scale collection of plant images and sensor data both in greenhouse and in field Chéné et al. (2012); Fahlgren et al. (2015); Hairmansis et al. (2014); Lin (2015); McCormick et al. (2016); Xiong et al. (2017). Figure 1 shows the field facility built by the Plant Science Institution (PSI) at Iowa State University, from which we can see that cameras are placing in front of each row of plants in a field. These cameras are designed to take photos with a certain frequency during the whole plant growth season. From the high-throughput system, we are able to process and extract useful phenotypical features, such as height, width and size, from the recorded images, and use those extracted features for plant genomics analysis. Compared to the traditional methods, the high-throughput system is able to provide the plant features of interest in a more efficient, accurate and non-destructive way.

**Figure 1.**
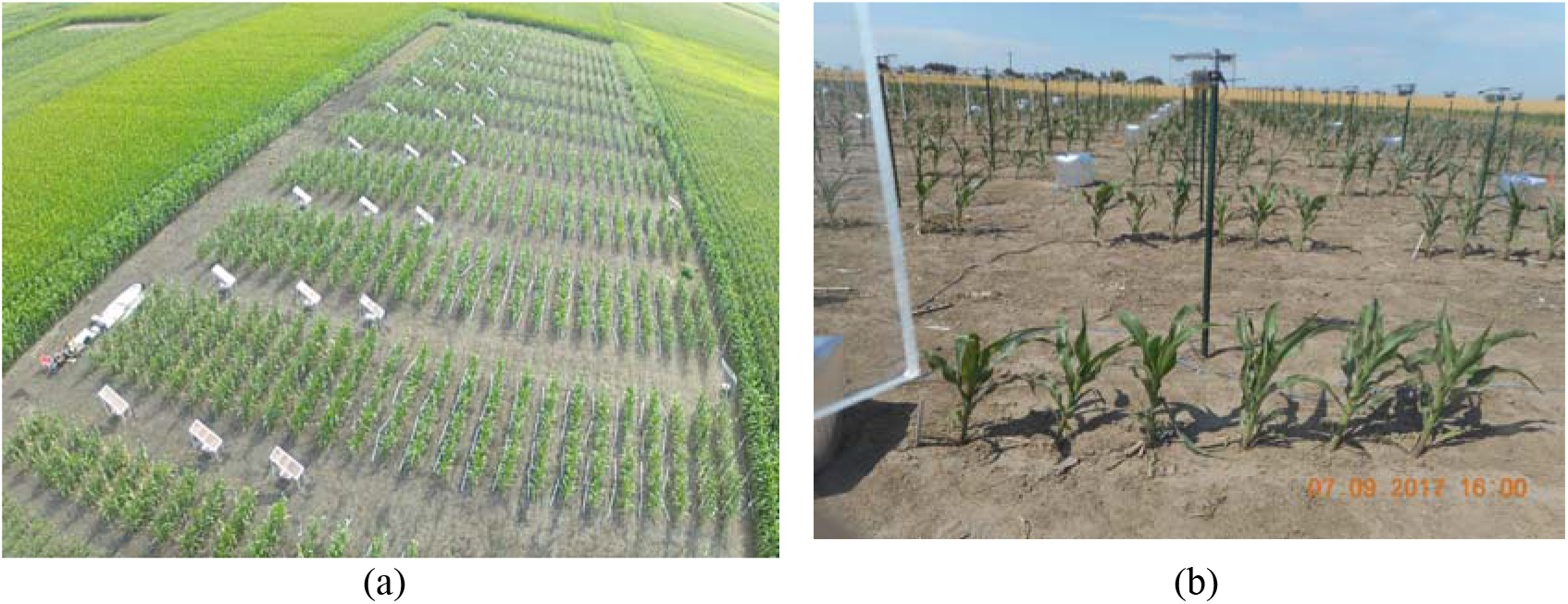
Left panel: an overview photo of the Iowa State field phenotyping system; right panel: raw RGB images of maize plants captured from the phenotyping facility.

Despite high-throughput systems can generate large amount of images per day, image processing is generally needed to extract numerical measurements of plant traits for downstream analyses (Adams et al., 2020; Choudhury et al., 2018). In order to extract phenotypical features from images, plant object segmentation is the fundamental step (Ge et al., 2016; Miao et al., 2018). However, image segmentation and traits extraction are the current bottlenecks in the area of high-throughput phenotyping. Separating plants from background is much easier for greenhouse images where the background is homogeneous (usually white) and a simple thresholding algorithm can provide satisfactory results (Choudhury et al., 2018; Ge et al., 2016). However, thresholding methods no longer works for field images as the backgrounds in the field are much more noisy than those in a well controlled greenhouse imaging chamber. See the background in Figure 1 as an example, which is a mixture of dirt and straws on the ground, pillar and white boxes as part of the phenotyping facility, and plant shadows.

Plant segmentation results in a binary image, where all pixels in the original RGB image are classified into either plant or background. Thresholding is the simplest and the most commonly used method for image segmentation (Davies, 2012; Hartmann et al., 2011), which classifies pixels by a cut-off value on pixel intensities. Thresholding could be applied on the average of red, green and blue channels, or the green-contrast intensity (Ge et al., 2016) obtained by green channel minus the average of blue and red channels, or both (Wang et al., 2020). Despite the popularity of thresholding methods for greenhouse images with uniform background color, they fail to work for images with noisy background. See Figure 2 as an illustration for the thresholding method on ISU field images of maize, where a smaller threshold (0.04) maintains most parts of the plants but leaves many background noises, and a larger threshold (0.08) removes most of the background noises but misses many pixels of the plants. Most importantly, the ideal threshold is sensitive to the environment and time of the image, which requires extensive human power for tuning.

**Figure 2.**
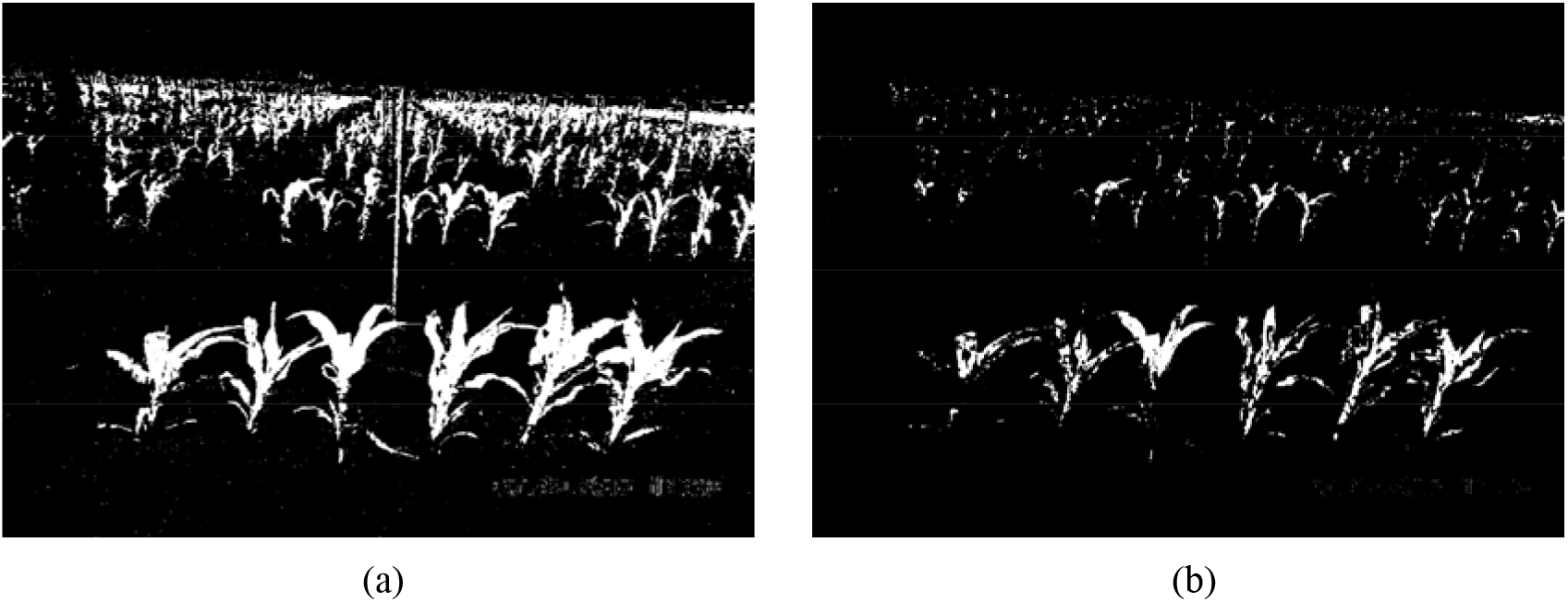
Thresholding segmentation method for Figure 1 by green-contrast intensity with weights 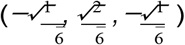, and threshold level 0.04 (left panel) and 0.08 (right panel).

A well separated plant image is the key to accurate feature extraction as traits like plant height and width are most sensitive to background noises. To improve thresholding methods for greenhouse images, Adams et al. (2020) made a thorough comparison for self-supervised learning methods trained on pixel intensities of plant RGB images from greenhouse, where the training data were obtained by unsupervised K-means clustering Johnson et al. (2002); Klukas et al. (2014). It is demonstrated that self-supervised neural network models are more accurate and robust than the traditional thresholding methods at segmentation. It is also worth to mention there have been an increasing number of applications of convolutional neural networks (CNN) to plant phenotype extraction from images in recent years. Miao et al. (2019) considered leaf counting of maize by a relatively shallow CNN; Lu et al. (2017) employed deep CNN structures to count the number of tassels on maize plants in field; Aich et al. (2018) used CNNs for estimating emergence and biomass of wheat plants.

In this paper, we develop an automatic and robust procedure to extract plant traits from field images generated by a high-throughput phenotyping system as shown in Figure 1, and fit an non-decreasing curve for the extracted traits over the plant growth period. The fitting is non-parametric and free of model assumptions, which can be used in any stage of plant growth. The first step of the proposed procedure is to obtain an accurate segmentation of plants from field images. Motivated by the idea in Adams et al. (2020), we construct a transform domain self-supervised neural network model, which use the plant pixels obtained by the K-means clustering algorithm from greenhouse images to train models for field images. The proposed method is built on neighborhood pixel intensities to separate the plant class from the background class for each pixel. We propose a novel self-supervised method to efficiently generate a large amount of training data for building the neural network model, which combines background pixels from field images and plant pixels from greenhouse images. The advantages of self-supervised learning are its easy implementation and efficient and automatic generation of data supervision, which avoids the time and labor intensive labelling process for preparing training data. Post-processing (Davies, 2012; Gehan et al., 2017; Hamuda et al., 2016; Vibhute and Bodhe, 2012) of the segmented image from the neural network model could be applied, such as median blur, erosion and dilation operations. Using the segmented images, we design a computationally efficient algorithm to identify and separate the target plants. Plant features can then be measured for each separated plant based on the segmented image. We also propose a refined feature extraction algorithm by pooling information of plant locations from a sequence of images taken over time in the same row of an experiment. In the last step, we fit a non-parametric and non-decreasing functional curve for the extract plant trait. The advantage of non-parametric functional fitting over parametric modeling and point-wise analysis of variance for plant growth dynamics are discussed in Xu et al. (2018). The proposed method restricts the fitted curve to be non-decreasing which leads to a more accurate estimation for growth curve comparing to the approach used in Xu et al. (2018). Comparing to the crowdsourcing height measurements by Amazon Turk workers, our method is automatic, more efficient and free of human labor. We also find that the proposed method produces more stable and consistent plant traits compared to the crowdsourcing results. Although we mainly focus on plant height measurement in this paper, the proposed procedure can be easily extended to extract other plant traits such as size and width.

## Method

The primary interest of this paper is to automatically extract heights of all the plants in the front row in Figure 1 for all the cameras in the field, and use the heights obtained in a sequence of photos taken over time to estimate plant growth curves. The proposed work flow from the original RGB image to the fitted growth curve for each plant is summarized in Figure 3. The main steps are enumerated in the following. The detailed procedures for each step are explained in the subsections.

1. Construct the train data set for plant and background pixels, where the plant pixels are obtained by K-Means clustering algorithm applied on plant images from greenhouse.
2. Perform image segmentation by neural network to transform the original RGB field image to a black and white image where white denotes the plants and black denotes backgrounds.
3. Identify the plants of interests and measure their heights from the segmented image.
4. Calculate the heights of plants from a sequence of images over the growth period.
5. Fit the plant growth curve using nonparametric regression with non-decreasing mean functions.

**Figure 3.**
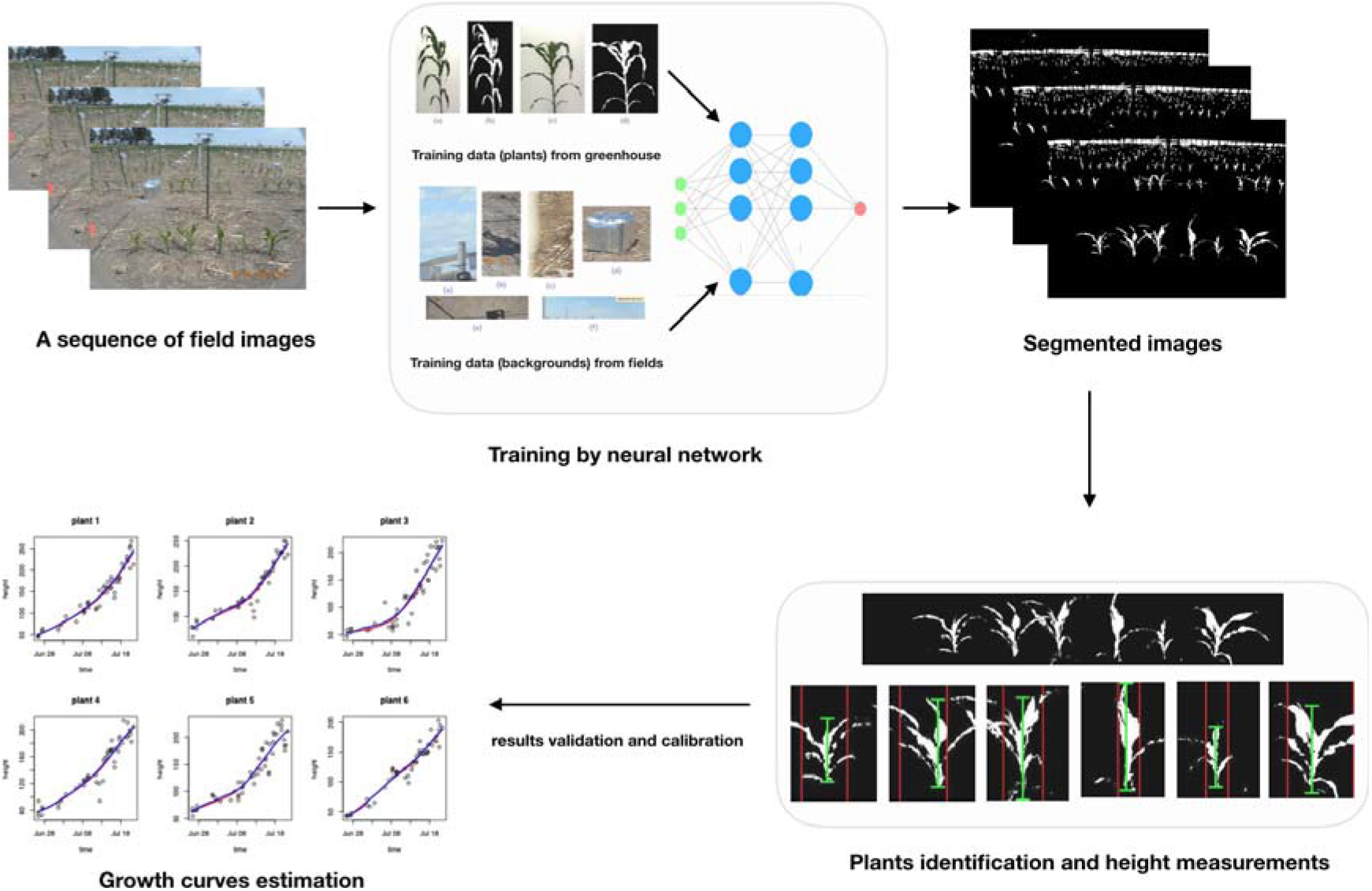
The diagram of the proposed method. From top left to bottom left are the algorithm workflow from the original RGB image to the fitted growth curves.

### Image data

In this paper, the image data we use were taken from a dry field in Nebraska in 2017. Two replications with 103 and 101 rows of maize plants were designed, where each row included six plants and one camera. The photos were taken with a frequency of 15 minutes, and the average number of photos taken by one camera is 1, 719 and 1, 650 respectively for the two replications. By only including the photo taken in the growth season, deleting problematic images (in dark environment or rainy and foggy weather, etc), and selecting the images taken around 10am everyday, there are on average a hundred of images per camera used in our analysis. For an illustration purpose, we randomly choose 10 cameras for each replication, and apply our algorithm to obtain the growth curves for all the plants taken by these 20 cameras. The raw field photos are high resolution (5152 *×*3864) RGB images with intensity values of red, green, and blue channels between 0 and 255 for each pixel. We normalize the pixel intensities by dividing 255, producing floating point numbers between 0 and 1. To increase the computation efficiency, we also re-scale the image resolution to 1000 *×* 750.

### Self-supervised learning

We consider self-supervised learning to classify each pixel of the field image into either plant class or background class. In supervised learning, collecting and preparing accurate training data is the most labor intensive and time consuming step. To overcome this major obstacle, we proposed an efficient self-supervised learning method to automatically construct training data with labeled pixels for field images. To prepare training data for background, it is straightforward to crop the image into pieces that only include the background. All the pixels in those pieces of images are labeled as background. For example, see the second panel in Figure 3, where the crops of background images include the dirt and straws on the ground, sky, shadows, and the phenotyping facilities.

To obtain training data for the plant class, however, it is time-consuming to accurately crop the field image to contain only the plants because of the irregular shapes of plants and noisy backgrounds in field images. We consider to borrow the plant pixels in greenhouse images, where plants are photoed in a well controlled imaging chamber, and the backgrounds are much less noisy than field images. Through cropping the greenhouse images, it is easy to obtain part of the plant under background with a universal color; see panels (a) and (c) in Figure 4 as an example. Motivated by the procedure proposed in Adams et al. (2020) for extracting plant pixels, we apply K-Means clustering algorithm with Euclidean distance metric on all the pixels in those cropped greenhouse images, and obtain the pixels in plant class; see panels (b) and (d) in Figure 4 as the clustering results from the original images in panels (a) and (c), respectively. All the extracted plant pixels from K-Means algorithm are collected as training samples of the plant class for the field images. The key idea is to use the pixels from greenhouse plant images to train the plant pixels in the field under the assumption that the intensities of the plant pixel are similar in the greenhouse and in the field. Note that there is no need to have a perfect segmentation of the whole plant from the greenhouse, as we only need part of the plant pixels where separation from the background is easy and can be done by K-Means clustering. Both the procedures construct training sample for the background and plant classes are easy to implement without human labeling and annotation. This makes the supervise learning for plant segmentation possible at the pixel level.

**Figure 4.**
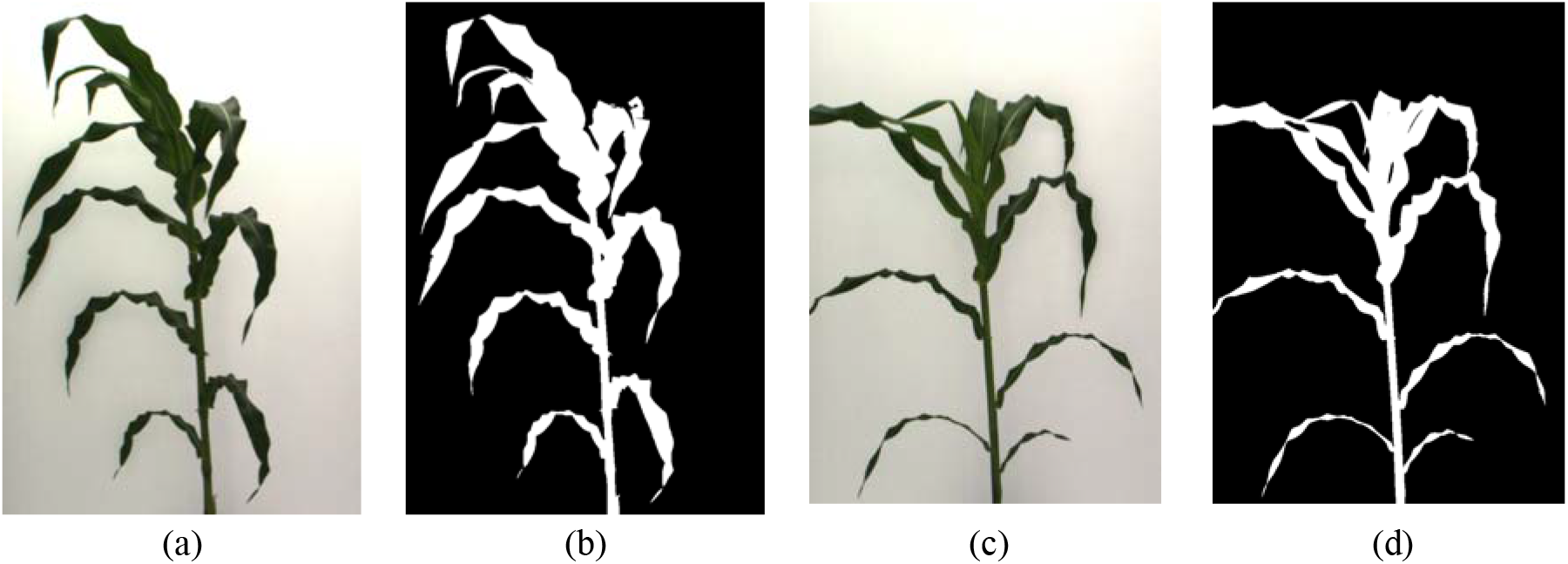
An example of training data (plant class) acquisition. Panels (a) and (c) are two cropped greenhouse images; panels (b) and (d) are the clustering results of K-Means algorithm (*K* = 3), the white parts are used as training data for the plant class.

### Segmentation by neural network

We use the training data containing 598,219 plant pixels and 2,728,415 of background pixels. For a given pixel, a 3 *×* 3 neighborhoods of that pixel together with their RGB intensities are used as the input features. This results in 27 features for each pixel. A three layer neural network under the API Keras in R is used to train the model. Specifically, the input layer has 27 nodes, and the first and second hidden layers have 1,024 and 512 neurons respectively. The ReLU activation function is used between the input layer and the first hidden layer as well as between the first and second hidden layers. The output layer had one neuron which gives the predicted probability of one particular pixel belonging to the plant class. The sigmoid activation function is used between the second hidden layer and the output layer. The dropout rates at each hidden layer are chosen to be 0.45 and 0.35 respectively. The binary cross-entropy loss function with the Adam optimization algorithm (learning rate = 0.001) is used to evaluate the network. Finally, we use 20 epochs with batch size 1,024 to train the model. 1% of the training data in each epoch are held out as validation data.

A cutoff threshold of 0.5 is used to classify the plant pixels, which means a pixel is classified as plant if its output probability from the neural net model is greater than 0.5. Figure 5 provides an example of the segmentation result by the proposed neural network model. We see that most of the plants are precisely segmented with few background noises.

**Figure 5.**
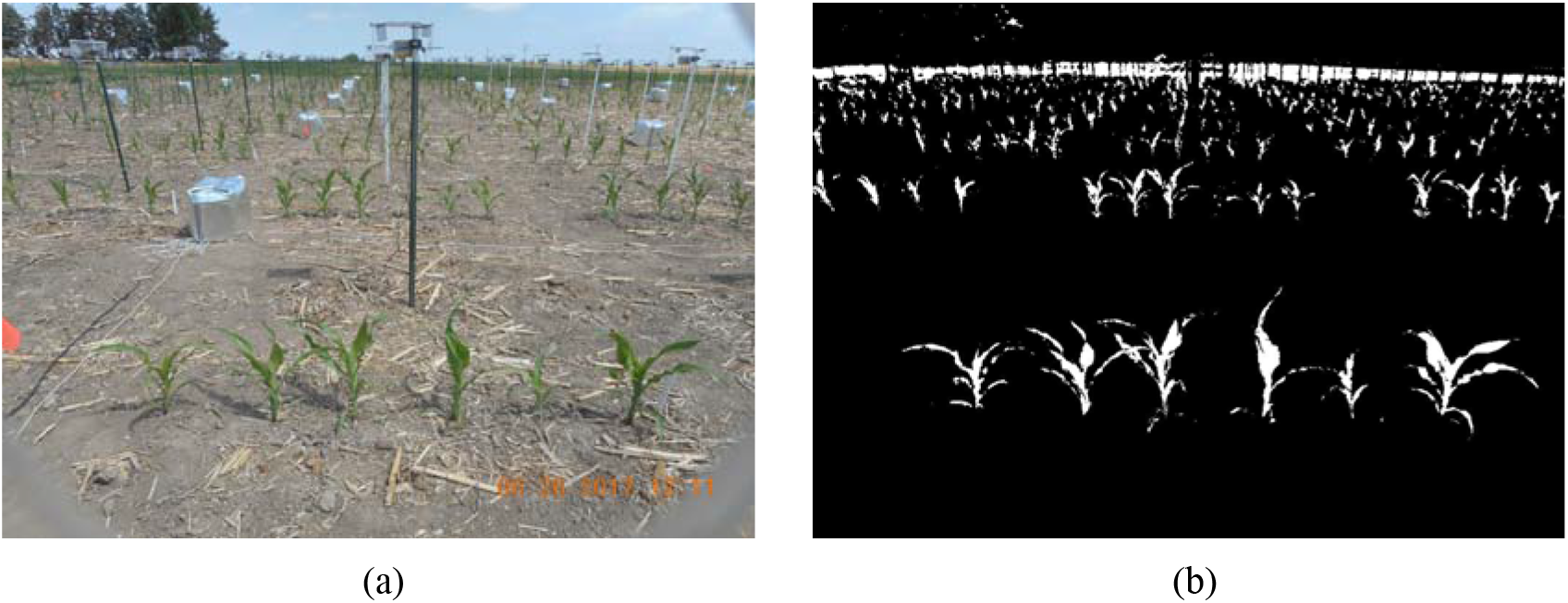
Segmentation result (right) of the original image (left) by the proposed self-supervised neural network model.

### Plant height measurement for a single segmented image

Based on the segmented images, we aim to measure the height of the plants in the first row of the image. As an example, there are six maize plants in the first row of Figure 5. This procedure constitutes identifying the first row by row cut, separating each plant in the first row by column cuts and measuring the individual height of each plant.

#### Row cut

To separate the first row in the image, we propose a row cut algorithm which consists of local maximum calling and region identification. Specifically, the row means are calculated for each row of the segmented image, which gives the percentage of plant pixels in each row. Then a local smoother (*loess* function in R) is used to smooth the row means. From Figure 6, we can see multiple peaks in the row mean curve, where the indexes of the bottom peak correspond to the plants in the first row. To find the local maximum of the bottom peak, we threshold the row means by *R*_*v*_ = 10% percent of their global maximum value. This results in segments of indices with value above the threshold, where two segments are considered as separate if they are *S*_*r*_ = 10 rows apart from each other. The maximal of the bottom peak is the largest row mean in the first segment from below. See the illustration in the top right panel of Figure 6, where the red point denotes the maximum of the bottom peak (colored in green) identified by the proposed procedure. Finally, to locate the region of the bottom peak, its upper and lower boundaries are chosen as the first rows smaller than *R*_*u*_ = 7.5% and *R*_*l*_ = 2.5% percentage of its peak maximum when moving away from the center of the bottom peak from above and below, respectively. See the bottom two panels in Figure 6 as an illustration for this step of region identification. Our results show that the proposed procedure can accurately separate the first row of plants and it is robust to the tuning parameters *R*_*v*_, *R*_*u*_, *R*_*l*_ and *S*_*r*_ for all the images we analyzed. However, their appropriate values may vary in a different setting of cameras.

**Figure 6.**
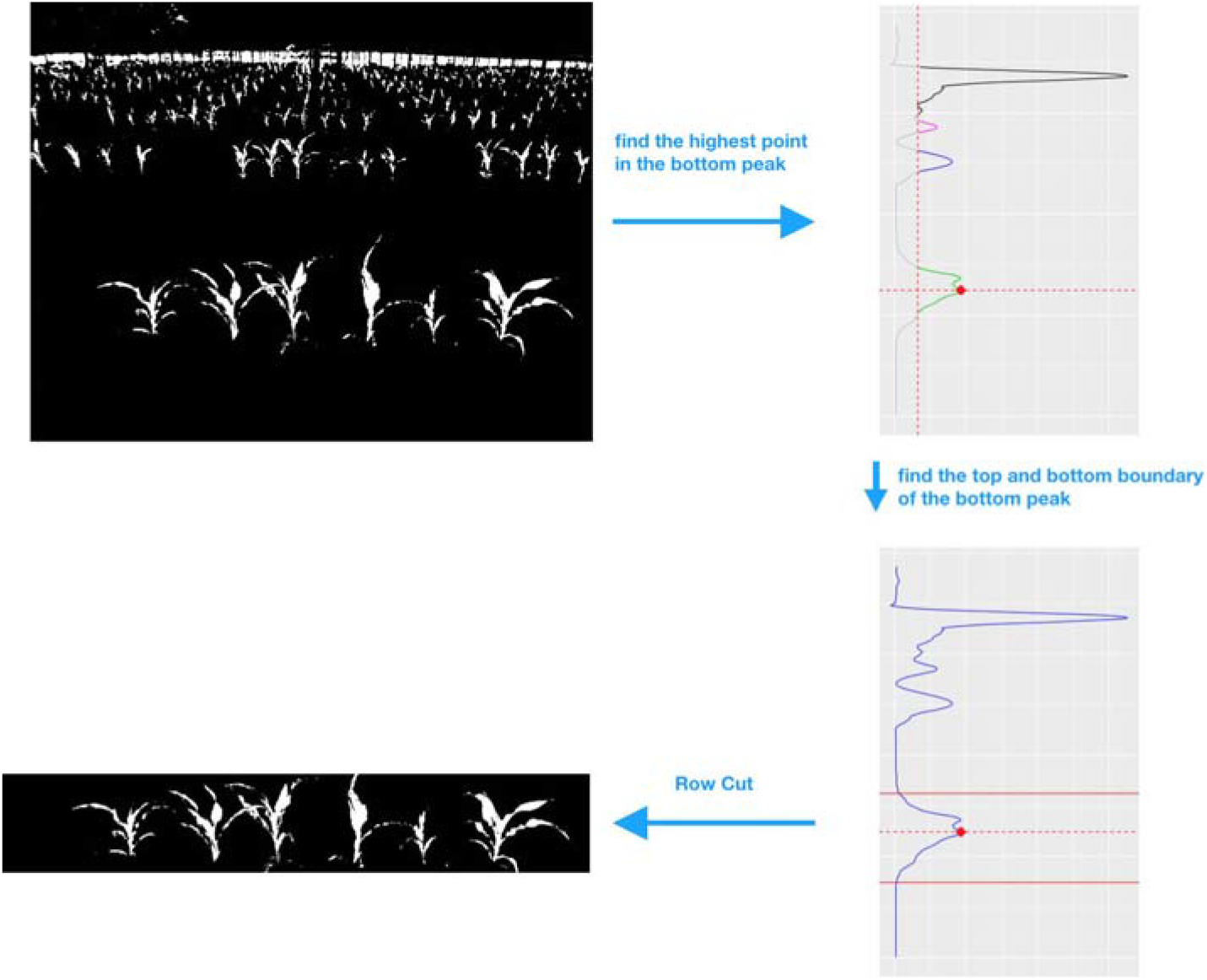
Diagram of the row cut algorithm. Top left panel: the segmented image of plants from the neural network model; top right panel: the step of local maximum calling, which provides a separation of different peaks (illustrated by different colors) in the row mean curve and find the maximum of the bottom peak (denoted by the red point); bottom right panel: the step of peak region identification, giving the upper and lower boundaries of the bottom peak (denoted by the red solid lines); bottom left panel: the segmented first row of plants from the original image.

#### Column cuts

Once the targeted row of plants is obtained, we separate each plant in that row by a column cuts algorithm. A diagram of this algorithm is shown in Figure 7 for illustration. Similar to the row cut algorithm, the first step is to compute the column mean values, which gives the column-wise percentage of the segmented plant pixels. We apply a quadratic power transformation (i.e. *f*(*x*) = *x*^2^) to the column means, which magnifies the column peak maximal values so that it is easier to separate different peaks, as illustrated in the third step in Figure 7. Following the same strategy as the row cut algorithm, we find the maximums for each peak by thresholding the squared column means at *C*_*h*_ = 20% percent of the overall maximum, and obtaining the column indexes with value larger than this threshold. Then, segments of indexes that are at least *S*_*c*_ = 50 columns apart are considered as from different peaks. The maximal value for each peak can be obtained as the largest squared column means in each segment. The cuts between plants are calculated as the middle points between the indexes of two adjacent peak maximums.

**Figure. 7.**
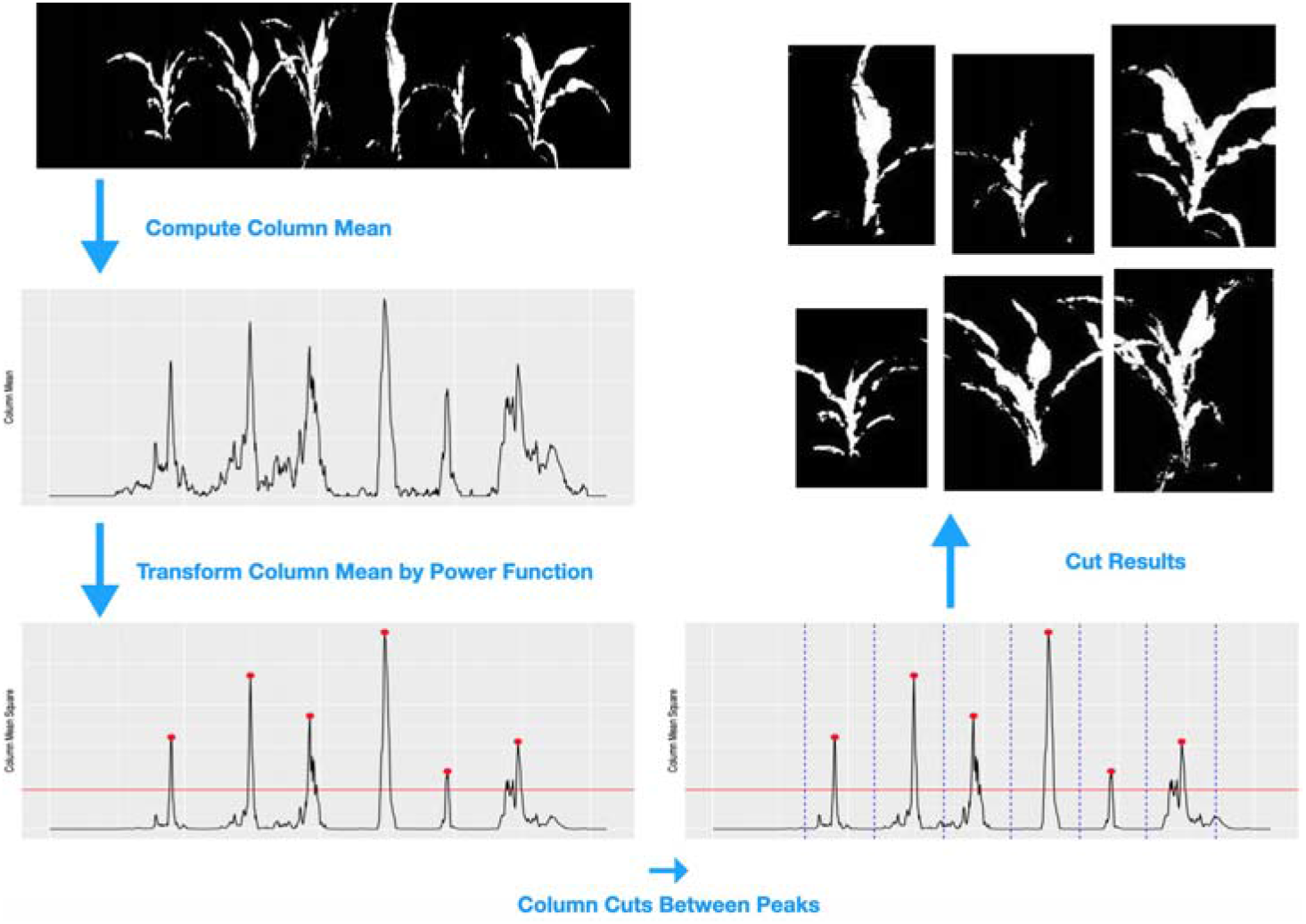
Diagram of the column cut algorithm. Top left panel: the segmented first row of plants from the row cut algorithm; middle left panel: visualize the plant distribution by the column mean curve; bottom left panel: the step of local maximum calling for the column mean curve, providing the maximum of each peak after the power transformation (denoted by red points); bottom right panel: the step of plant separation, where the cuts (blue dashes lines) between plants are calculated as the middle points of two adjacent peaks; top right panel: the segmented image for each plant.

Specifically, let 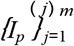 be the indexes of the column-mean peak maximum for the *m* plants. The indexes of the column cuts are 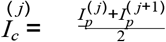 for *j* = 2,....*m*. The left and right margin cuts are defined to be 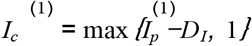 and 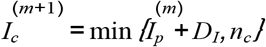 respectively, where 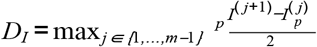 and *n*_*c*_ is the total number of columns.

#### Phenotype measurements

After making the row and column cuts, we can measure phenotypic traits for each plant. In this study, we focus on height measurement. The proposed procedure can be easily adjusted to calculate plant width and size. For the height of each separated plant, we first compute the column means, find their maximum value and the corresponding index of the maximum. Then, the left and right cuts are made to retain the center part of the plant, where the two cuts are chosen as the closest columns to the highest column that are smaller than 10% of the maximum value. The row mean values for the selected center part of the plant are computed, and the plant height is calculated as the index difference between the first row from below and the first row from above with mean values larger than 2.5% of the maximal row mean value. The diagram of the proposed procedure for height measurement is shown in Figure 8.

**Figure 8.**
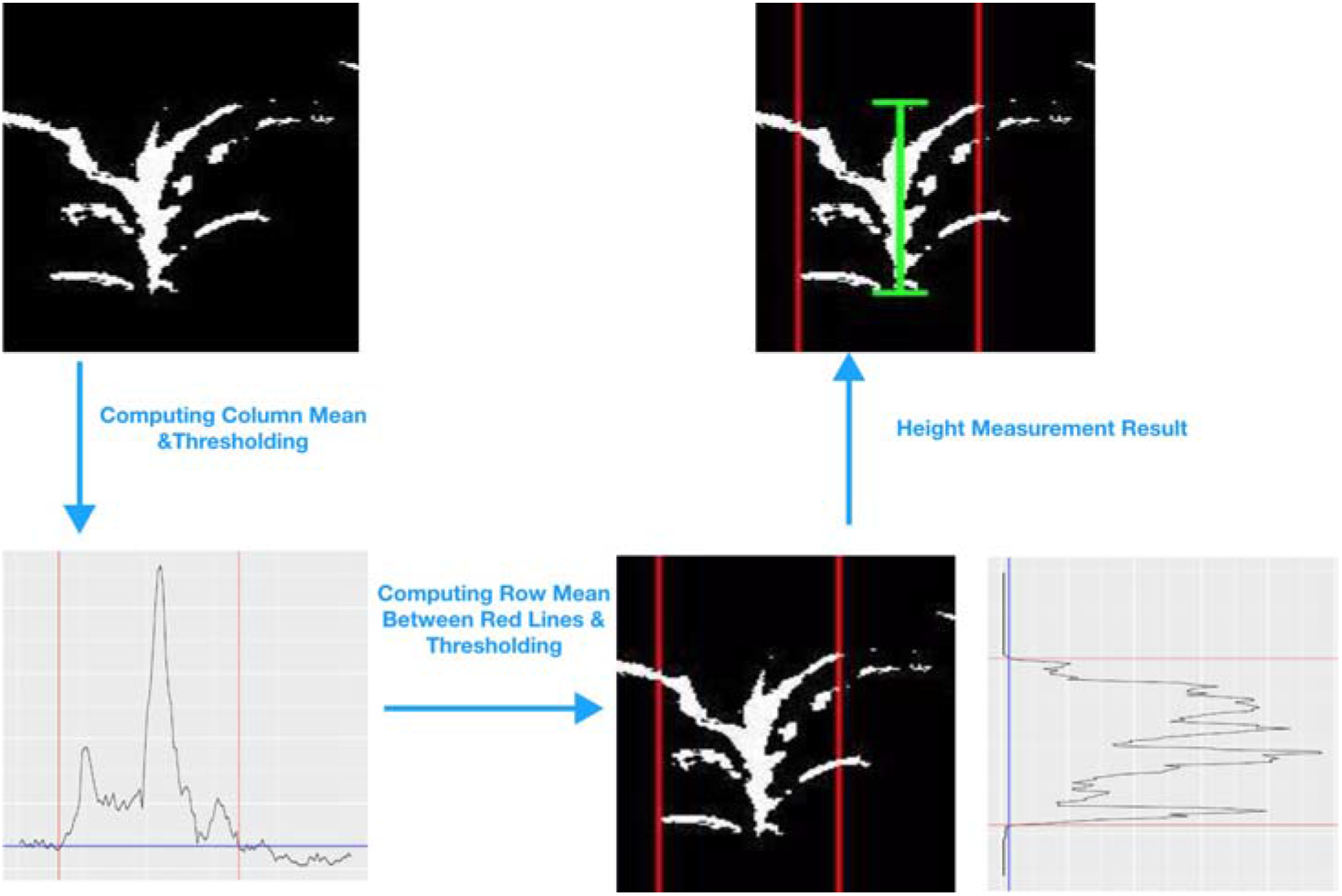
Diagram of the height measurement algorithm. Top left panel: the segmented image for a single plant from the row cut and column cuts algorithms; bottom left panel: extracting the center part of the plant by thresholding (blue line) the column mean curve of the segmented image in the top left panel and identifying the left and right cuts (red lines); bottom right panel: the extracted center part (marked by two solid red lines) of the segmented image, and the height measurement by thresholding (blue line) the row mean curve of the center part of the segmented image; top right panel: the segmented image of a plant with the annotated height.

### Plant height measurement for time series of images

In this section, we propose a refined height measurement procedure for a sequence of plant photos taken over time by borrowing information of plant locations across the time series of images. After conducting the above procedures for image segmentation, row cut and column cuts, we can systematically study the growth trend of each separated plant over time, and refine the column cuts algorithm that is solely based on one image by considering a sequence of images from the same row, as the camera positions are roughly kept fixed during the experiment. This can also help to remove the problematic images and images with overlapping rows of plants from which a clear separation of the plants in the front row is difficult.

Figure 9 shows a set of field photos from the same row of plants taken by the same camera over time. Notice that the location of those plants are roughly kept the same across different photos. However, we can not identify all the six plants from every photo due to technique issue of the camera (panels a and b where the plant on the most right side is outside of vision), strong wind (panel e where the second and third plants overlap with each other) or death of plants. Meanwhile, the row cut algorithm requires a separation between the first row and the second row of plants, so that the bottom peak of the row means are separable from other peaks; see Figure 6. When the plants in the first row overlap with the plants in the background, as shown in panel (f) of Figure 9, it is challenging to accurately measure plant heights by computer vision methods. The proposed method is designed for the earlier growth stage of plants in fields. Our algorithm of neural network is not able to separate the first row from the rest of rows if they are overlapping. We will discuss possible solutions for this problem in the discussion section.

**Figure 9.**
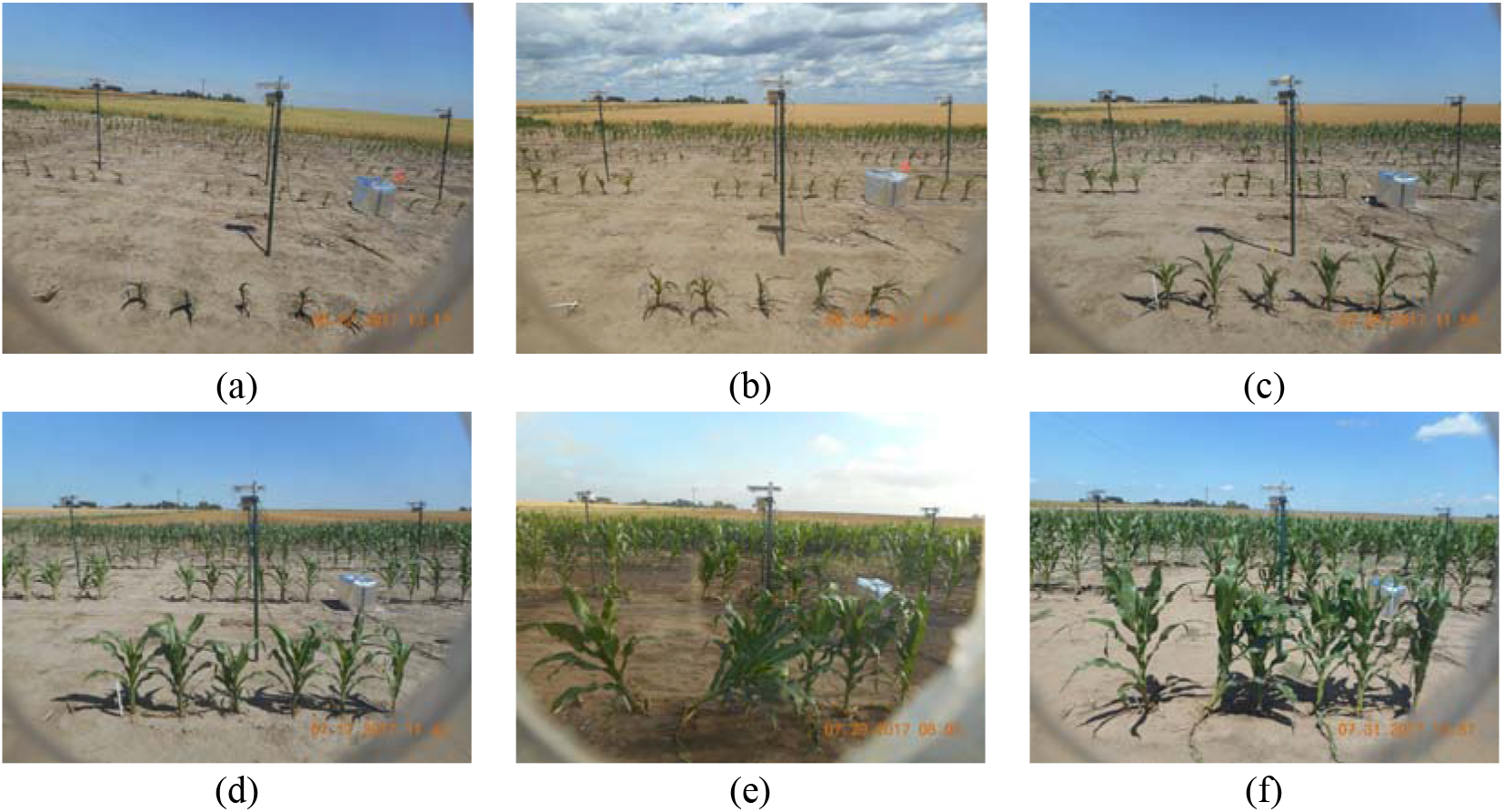
A sequence of field photos from the same row of plants over the growth period.

To deal with the aforementioned challenges of the dynamic photos of plant growth, we propose method to check quality of the images in order to obtain reliable plant height estimation. The algorithm includes four steps as follows. Firstly, the neural network segmentation model and the row cut algorithm are applied to every photo in the sequence, and the heights of the segmented first row from each image are computed. We apply change point detection methods (via *changepoint* R package) to identify jumps in the heights of the segmented rows from the sequence of images. As illustrated in the top left panel of Figure 10, there is a clear jump in the row heights around July 21. This change point, denoted by the red vertical line, corresponds to the date when the front line of plants overlap with the plants in the background, and become inseparable. We focus on measuring the plant heights of the front row before the change point. Secondly, column cuts algorithm is implemented to count the number of plants in the front row for the segmented images from step one. The mode of these counts, denoted by *m*, is used as an estimate for the true number of plants in a given row over time. As six seeds are planted in each row in this experiment, the modes for most of the rows are six in the growth period. We only consider the images with the number of plants in the first row equal to its mode *m*. This is illustrated in the top right and bottom left panels of Figure 10, where *m* = 6 and the red points are the images with 6 identified plants over the time course. We compute the plant heights for those selected images for the time sequence of photos in the following steps.

**Figure 10.**
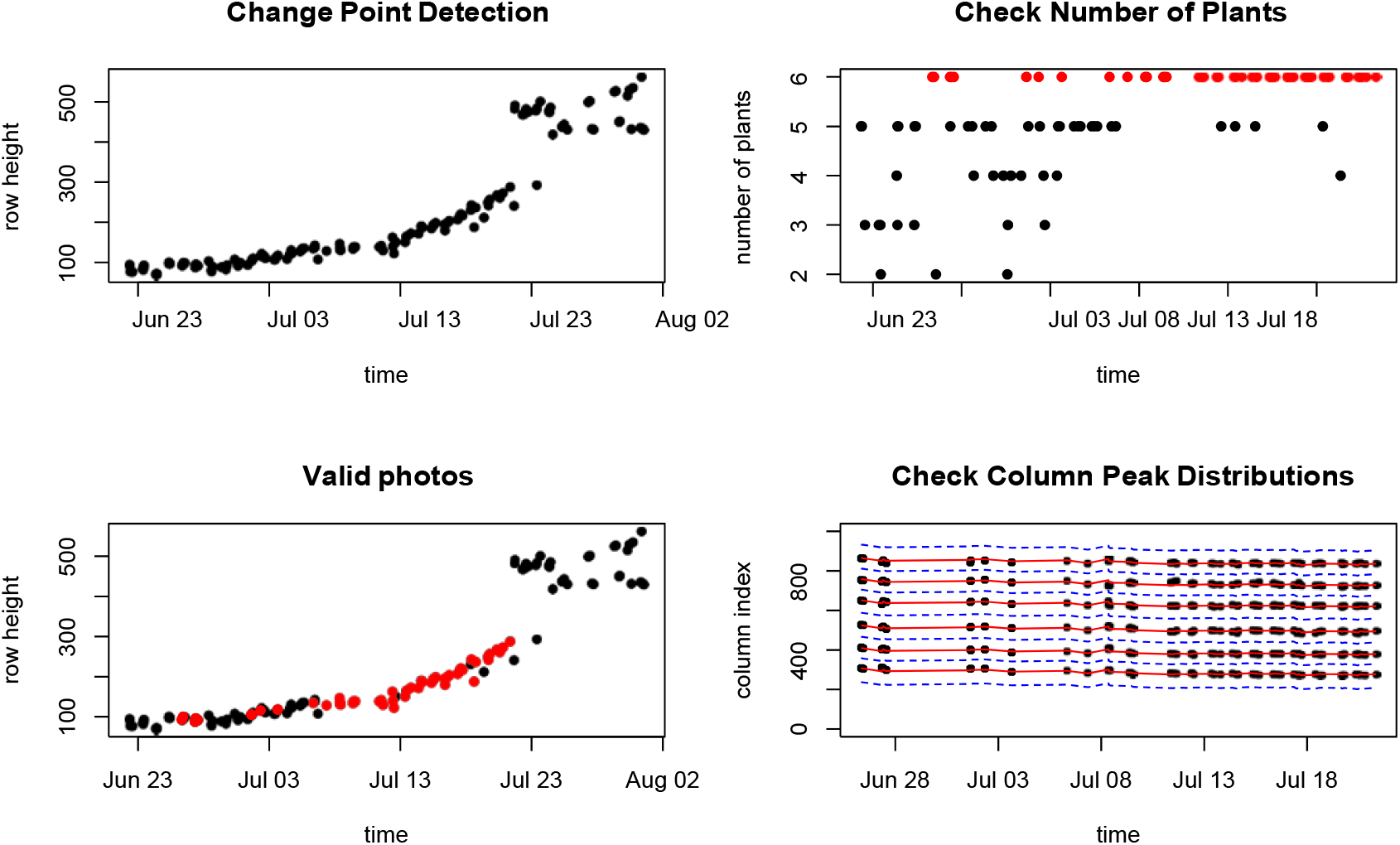
Refine the height measurements for a sequence of images. Top left panel: change point detection to identify the jump in the heights of the segmented rows, where the plants in the first row overlap with the background rows; top right panel: the number of identified plants in a given row over time, where 6 is the mode; bottom left: the selected images (marked as red) for the growth curve analysis, which have 6 identified plants before row overlapping; bottom right: refining the column cuts for each image by pooling information of plants location from other images in the same row over the growth period, where the red solid lines are the estimated center of each plant over time, and the blue dashed lines are the refined column cuts.

Given a row (camera), let *n* be the number of the selected images with *m* identified plants from the first two steps. In the third step, we refine the column cuts for each plant in a row by pooling information of plant locations from those selected *n* images. Let 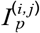 be the column peak index for the *j*th plant in the *i*th photo. The average column peak index for the *j*th plant can be computed as 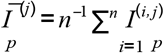. Note that the camera might slightly shift horizontally, which affects the position of the column peaks over time in a given row. However, the distance between two adjacent peaks should hold the same. Therefore, it is reasonable to stabilize the column peak index for the *j*th plant in the *i*th photo as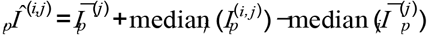, where the term 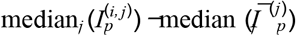 adjusts the horizontal shift of the camera. The separation for each plant can be made at the average index of two adjacent peaks, as discussed in the “Column cuts” section. The red solid lines and blue dashed lines in the bottom right panel of Figure 10 show the stabilized column peaks and column cuts, respectively. Finally, given each separated plants, we calculate their heights as discussed in the previous section. The measured heights for the six plants in Figure 10 are shown in Panel (a) of Figure 11.

**Figure 11.**
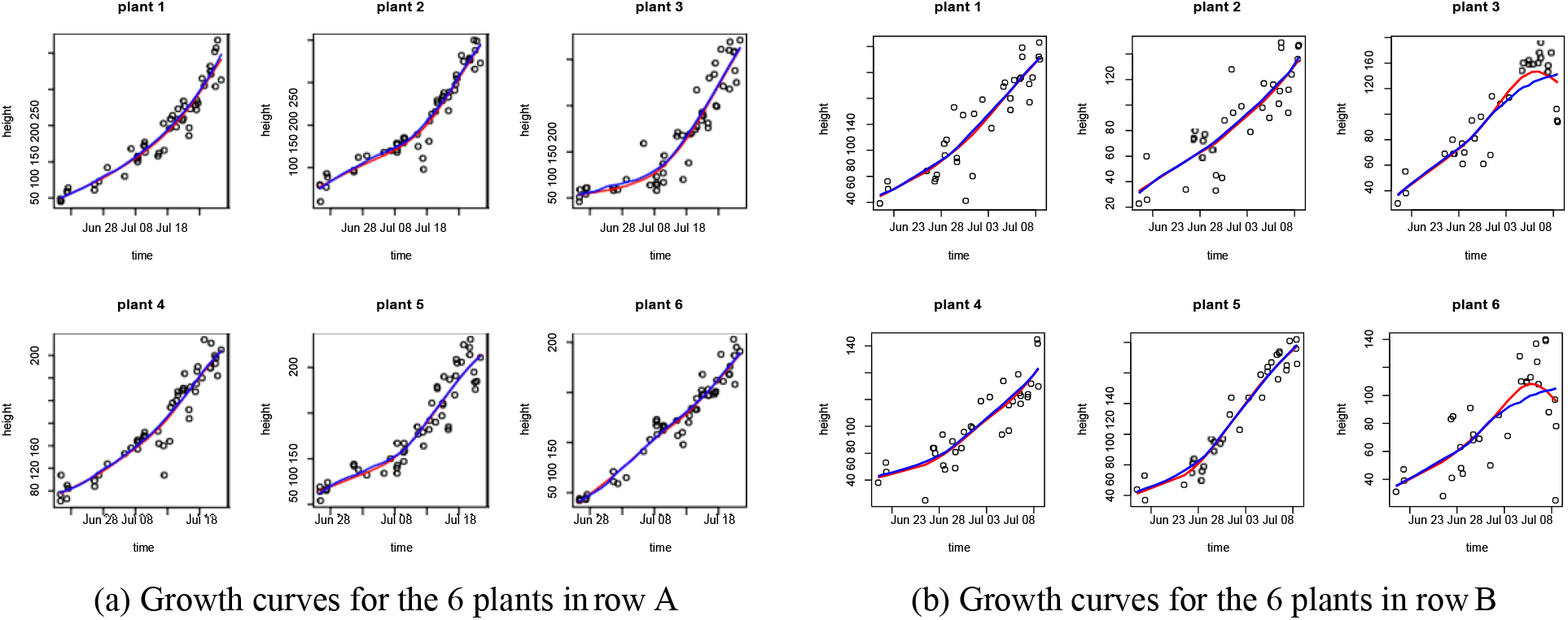
The fitted growth curve plots for each plant in two sets of images from two different rows. The points are the extracted plant heights from images; the red curves denote the classic penalized spline regression fit, where non-decreasing property is not guaranteed; the blue curves denote the non-decreasing fit we apply in this paper.

### Growth curves estimation

Based on the extracted heights from the plant images, we can fit a growth curve for each plant by nonparametric regression (Fan and Gijbels, 1996; Wahba, 1990). The red curves shown in Figure 11 are the smoothing spline fit for the plant heights. We can see that smoothing spline can capture the growth pattern well for most of the plants, however, it cannot ensure the non-decreasing property for the growth curve, as shown for plants 3 and 6 in panel (b) of Figure 11. To fit a non-decreasing function for the plant growth, following Dette et al. (2006), we first apply a kernel based estimation to fit an unconstrained growth curve 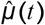. Then, we construct a density estimate using the estimated values 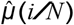 for *i* = 1*, ..., N*, where *N* is the total number of observations over time. It can be shown that integrating the density estimate from *−*∞ to *t* gives a consistent and non-decreasing estimator for *μ*^−1^(*t*) if *μ*(*t*) is a non-decreasing function. Thus, the estimator for *μ*(*t*) is also a non-decreasing function. The blue curves in Figure 11 are the fitted non-decreasing growth curves based on this method. We can see that the monotone fitting method solves the decreasing pattern of the estimated growth curve for plants 3 and 6 in panel (b) due to the high variation of measurements near the end points of this study. Meanwhile, the monotone fitting results are almost identical to the results by smoothing splines for other plants.

## Results

In this section, we compare the plant heights computed by our proposed method to the crowdsourcing height measurements by Amazon Turk workers. Figure 12 gives one example of these two measurements. The annotated image in panel (a) illustrates the heights obtained by the proposed method, where the red horizontal and vertical lines denote the row and column cuts, and the green vertical lines draws the heights at the center of each plant. Panel (b) in Figure 12 gives the bounding boxes for each of the plants draw by paid Amazon Turks, where the height is calculated as the difference between the top and bottom edges of the bounding box. It is clear to see that several of those bounding boxes are much higher than the plant, and some of them do not cover the entire plant. This happens frequently in the crowdsourcing results. The measurements based on Amazon Turks may lead to inaccurate and over-estimated plant heights.

**Figure 12.**
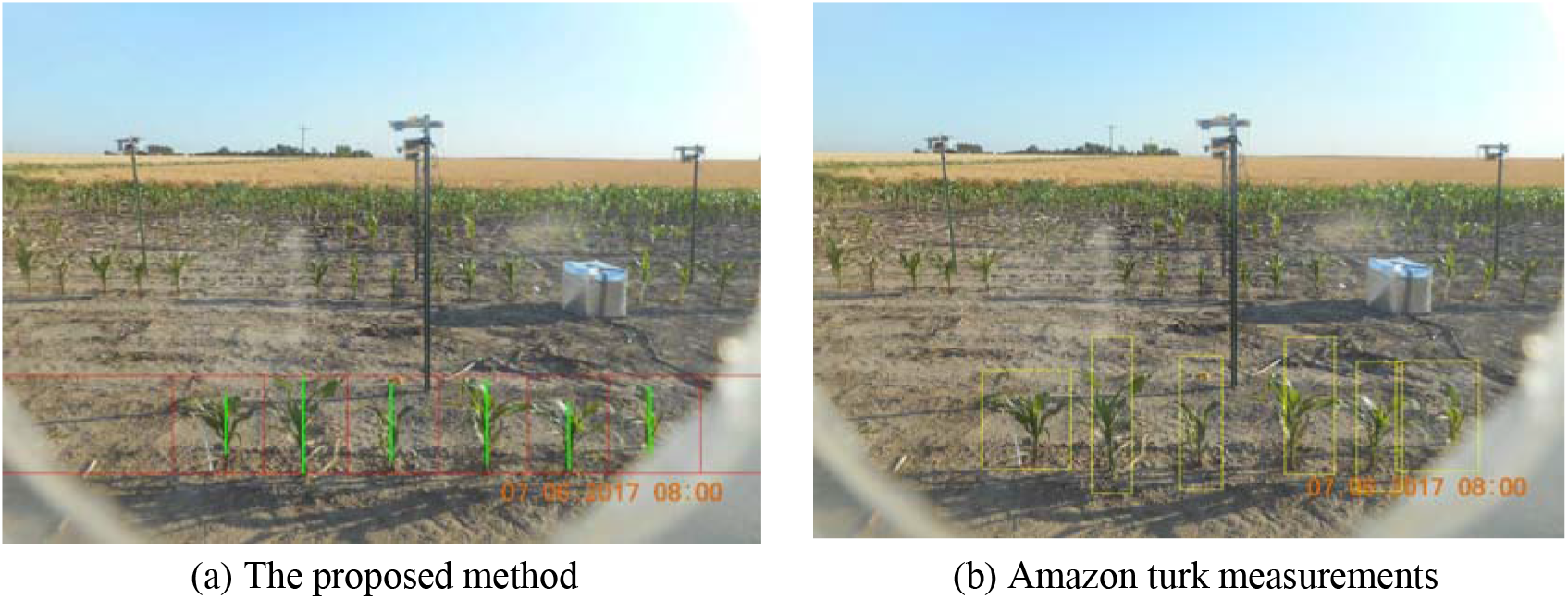
Visual comparison between our method and Amazon Turks measurements.

In panel (a) of Figure 13, we plot a sequence of crowdsourcing height measurements for the 2nd plant (from left to right) in Figure 12. From Figure 13, we can see high variability of the crowdsourcing measurements, where each image is annotated by three Amazon Turk workers. Panels (b) and (c) in Figure 13 respectively show the average height for each image over the three workers and the heights automatically calculated by the proposed method. Compared with panel (c), we can see that the Amazon Turks provide a less robust results with higher variation, even after averaging the repeated measures from different workers. The proposed method can provide better measurements for plant traits due to the accurate segmentation of plants by the neural network model. Panel (d) in Figure 13 provides a comparison for the two methods on plant height, where the red dashed line denotes the 45 degree line, and the red solid line is the linear regression fitted line. We can see that the results produced by the proposed method and Amazon Turk have a high correlation with *R*^2^ = 0.8119. But, our method consistently provides relatively smaller measurements for heights, as the bounding boxes tend to over-estimate the plant heights as illustrated in Figure 12.

**Figure 13.**
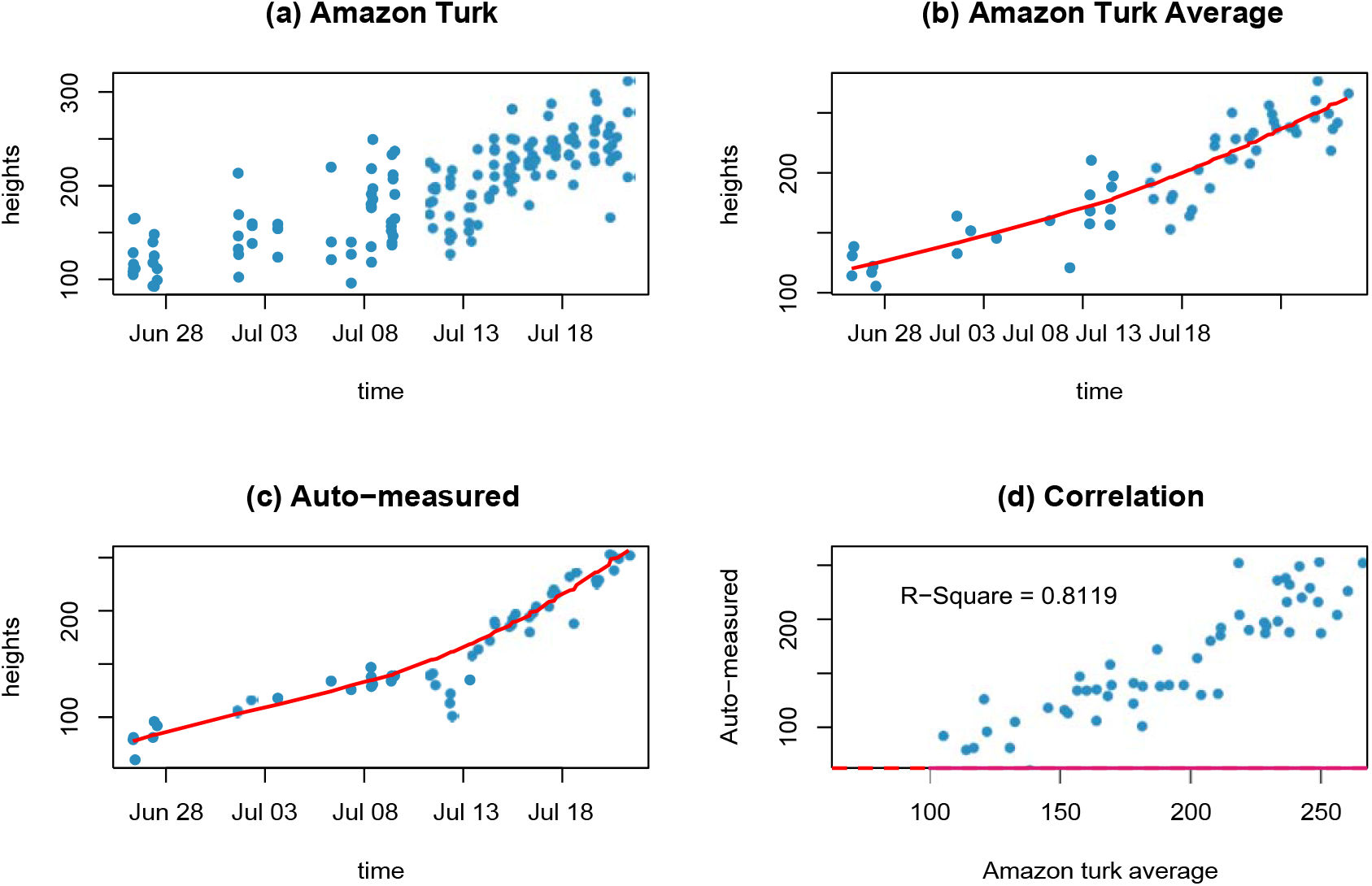
Comparison of height measurements (for one example plant) between the proposed method and Amazon Turks (crowdsourcing). Panel (a): crowdsourcing measured heights with each image annotated by 3 workers; (b) average heights from Amazon Turks with the fitted growth curve; (c) heights calculated by the proposed method with the fitted growth curve; **(d)** The scatter plot of heights measured by the proposed method versus Amazon Turk averages.

To further illustrate the proposed method lead to a more stable estimation for the growth curve, we compare the sum of square error (SSE) of the fitted curves of plant height between the proposed method and the crowdsourcing measurements. We calculate the ratios of the SSE from the crowdsourcing heights over that from the proposed method. The boxplots of the SSE ratios are presented in Figure 14, where panels (a) and (b) give the SSE ratios for 10 cameras (rows) and six plant positions, respectively. We can see that the proposed method provide a smaller SSE on average for each camera and each plant position, as most of the ratios are higher than 1.

**Figure 14.**
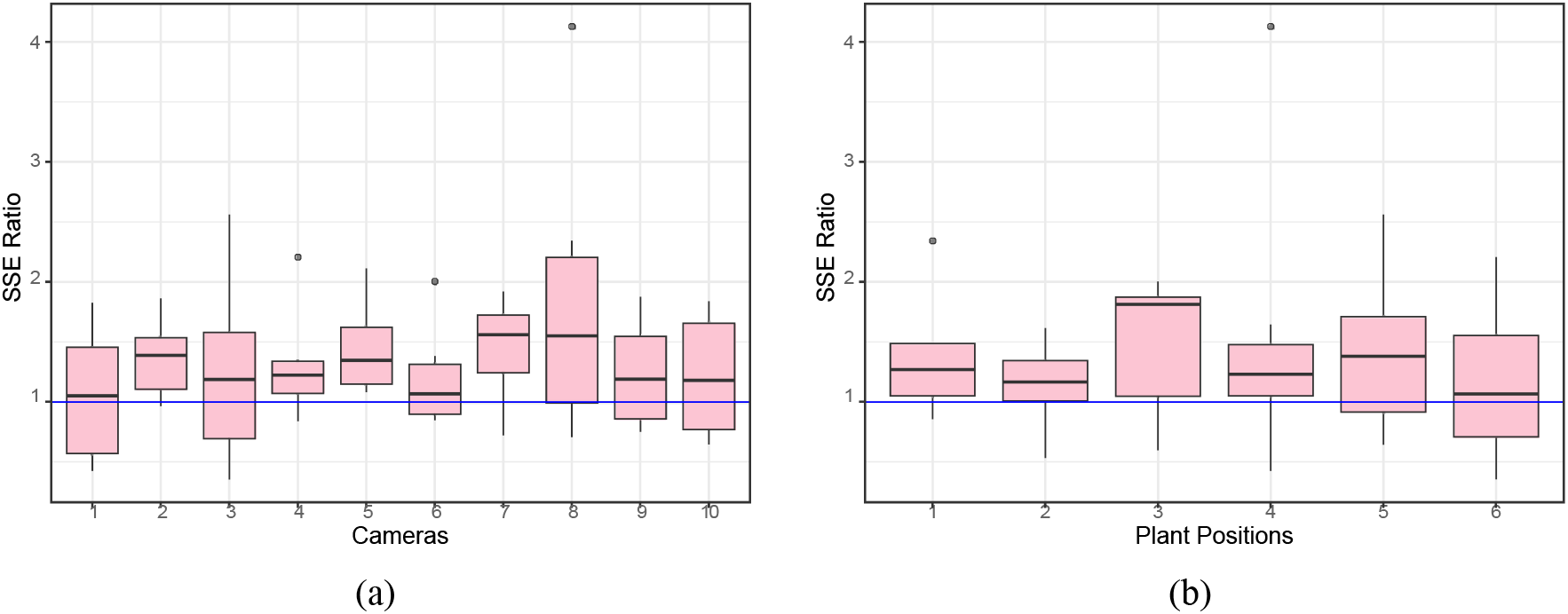
The SSE ratio of height measurements (for 10 cameras 6 plants) of Amazon Turks over the proposed method: (a) the boxplot of SSE ratios for 10 randomly select cameras; (b) the boxplot of SSE ratios for the six plant positions.

## Discussion

This paper provides a self-supervised method to separate plants from background for field images and a computation pipeline to extract plant features from the segmented images. Self-supervised learning is advantageous for high-throughput phenotyping as no human-labelling is needed to construct supervisory training data. This makes the proposed method easy to implement and broadens its application in plant phenotyping. The idea of transform learning that uses greenhouse images to learn field images can be applied in various feature extraction problems. As many plant features, including height and number of leaves, have been extracted from greenhouse plant images (Miao et al., 2019), we can generate pseudo field images based on greenhouse images with their extracted plant features, and build machine learning models on those pseudo images to measure plant traits for field phenotyping facilities.

As shown in Figure 10, the proposed method works for early stages of plant growth, where the first row in the images does not overlap with plants in background. Self-supervised learning methods can also be developed to separate the first row from the background plants if they overlap. This can be achieved in a two-step procedure. In the first step, the proposed segmentation method is applied to segment all plants from background. Training data of plant pixels from the first row and the background rows can be automatically formed from the images where the first row is separable. In the second step, using the training data, a CNN model can be constructed based on the pixel intensities from a small neighborhood of each pixel. This idea is to use plant images in early growth stages to form self-supervisory for separation of plants in late growth stages.

The proposed functional curve smoothing method is applied on each individual plant over time. Functional data analysis for genotype and treatment effects on plant growth can be conducted based on the fitted values from the non-decreasing functional curve. The “implant” package (Wang et al., 2020) can be applied on the smoothed plant traits for this purpose.

## Additional information

The R codes of the proposed pipeline, sample image data and description are available on Github at https://github.com/xingcheg/Plant-Traits-Extraction

## Author contributions statement

X.G., Y.Q. and D.N. developed the pipeline, conducted the analysis, and wrote the manuscript; C.T.Y., Z.Z., S.H. and P.S.S. conceived the experiment; C.T.Y. managed image data storage and access.

